# DistPCA: Tera-Scale Genomic PCA via Out-of-Core Distributed Parallelism

**DOI:** 10.64898/2026.05.15.725487

**Authors:** Georgios Mermigkis, Argiris Sofotasios, Eugenia-Maria Kontopoulou, Efstratios Gallopoulos, Panagiotis Hadjidoukas

## Abstract

Principal Component Analysis (PCA) is a fundamental tool in human genetics, widely used to study population structure. However, the rapid growth of modern genomic datasets, which often exceed main memory capacity, renders traditional PCA methods infeasible, motivating out-of-core approaches. Prior work on out-of-core genomic PCA has focused primarily on optimizing the inherently compute-intensive numerical core, largely overlooking the stages of data I/O and preprocessing, which emerge as significant performance bottlenecks at tera-scale. Furthermore, existing approaches remain limited to shared-memory single-node architectures, lacking support for distributed multi-node environments. To address these limitations, we introduce DistPCA, the first distributed out-of-core framework for tera-scale genomic PCA, implemented as a C++ library and scalable across both single- and multi-node systems. Built on top of Message Passage Interface (MPI), the proposed framework employs multi-level data parallelism across the entire PCA pipeline, combining multiprocessing, multithreading, SIMD vectorization, and compute–transfer overlap, while remaining compatible with block-based methods that rely on associative operations. Extensive evaluation on real and synthetic datasets demonstrates near-linear scalability, achieving speedups of up to 58.2× and over 98% reduction in wall-clock time, while maintaining parallel efficiency above 82% and preserving accuracy in the recovered principal components.

## I. Introduction

**P**rincipal Component Analysis (PCA) is a fundamental statistical technique for unsupervised linear dimensionality reduction. Originally introduced by Pearson [1], reformulated by Hotelling [2] and formally analyzed and systemized by Jolliffe [3], PCA identifies an ordered set of orthogonal directions, referred to as principal components (PCs), that capture the maximum variance of the input data. Due to its ability to summarize high-dimensional data through a smaller feature space of uncorrelated and informative variables, PCA has become a cornerstone tool for exploratory data analysis across a wide range of scientific domains [4]–[7].

One such domain is bioinformatics, where input biomedical data are typically high-dimensional and composed of a large number of interrelated variables, making PCA a standard tool for their analysis and interpretation [8]. Common applications include dimensionality reduction and clustering in gene expression studies [8]–[10], denoising and visualization of microarray data [11]–[13], exploratory analysis in metabolomics data [14]–[16], and analysis of molecular dynamics simulations of biomolecules [17]. PCA is also widely used in single-cell genomics, where it serves as an initial preprocessing step for downstream analysis tasks such as clustering, visualization, and batch-effect correction [18]–[20]. Finally, in population genetic analyses, PCA is employed to identify and characterize individuals and populations from genotype matrices, and to infer historical and evolutionary conclusions regarding their origins, migration, and genetic relatedness [21]–[24]. Seminal studies have demonstrated its ability to recover geographic patterns of genetic variation [25], [26], establishing it as a commonly used technique for ancestry inference and population stratification correction in Genome-Wide Association Studies (GWAS) [27]–[29].

Over the past two decades, the scale of genomic datasets has increased dramatically [30], [31]. Early efforts, such as the 1000 Genomes Project [32], included thousands of individuals, whereas modern biobanks, such as the UK Biobank [33], now provide sequencing data for hundreds of thousands of samples. These datasets enable large-scale studies of rare variants and complex traits, but also introduce significant computational challenges that push the boundaries of what conventional single-node systems can handle.

To accommodate this scale, genotype data is typically stored in compressed formats such as PLINK .*bed* files, where each genotype is encoded using a compact two-bit representation [34]. While this approach significantly reduces storage demands, PCA computation requires access to decompressed normalized representations of the data in main memory (RAM). For modern datasets, which expand to terabytes when decompressed, this requirement exceeds the RAM capacity of typical systems, making in-core computation infeasible [24], [35]. As a result, recent approaches have adopted out-of-core strategies that process the data incrementally in blocks streamed from storage [36]–[38].

Out-of-core PCA pipeline is inherently decomposed into three stages: (i) the *I/O* stage, responsible for fetching data blocks from storage, (ii) the *preprocessing* stage, which performs decoding and normalization, and (iii) the *numerical method* stage, which computes the sought PCs. The first two stages form the *input data pipeline*, which prepares the data blocks for the subsequent linear algebraic operations in the third stage. Empirical studies have shown that, in current out-of-core PCA approaches, the input data pipeline accounts for the majority of execution time, often exceeding 50% and reaching up to 75% of total wall-clock time [36], [38]. These findings indicate that, at tera-scale, the performance of PCA is no longer dominated by numerical computation, but instead by the cost of data movement (i.e., I/O operations) and preprocessing.

Nonetheless, most existing approaches for large-scale genomic PCA focus primarily on accelerating the numerical method stage through optimized linear algebra kernels and shared-memory parallelism [36]–[39], while the input data pipeline stages remain sequential. This introduces two key limitations: (i) confinement to single-node environments, and (ii) neglect of the performance overhead introduced by repeated I/O operations for data fetching and preprocessing. These limitations become increasingly restrictive as dataset sizes continue to grow, preventing effective utilization of distributed computing resources and leading to scalability challenges.

Motivated by these observations, we propose **DistPCA**, a novel distributed out-of-core framework for scalable genomic PCA that shifts the performance optimization focus from the underlying numerical method to the entire data processing pipeline, while preserving the accuracy of the recovered PCs. Built on top of Message Passage Interface (MPI), our framework can be deployed on both private and public cluster/cloud infrastructure, supporting efficient I/O performance on parallel file systems (e.g., GPFS), while leveraging high-performance network communication in a portable manner. The main contributions of this work are as follows:

- A hybrid multi-level parallelism scheme that combines MPI-based distributed execution with OpenMP multi-threading, Single Instruction Multiple Data (SIMD) vectorization, and double-buffered I/O to overlap computation and data transfer across all stages of the pipeline.
- A holistic approach that aims to jointly optimize the three key stages of out-of-core PCA, supporting efficient and scalable execution across multi-node High-Performance Computing (HPC) clusters and achieving performance gains of up to 5.26× over prior solutions that remain confined to single-node environments.
- A method-agnostic design that allows seamless integration with block-based PCA algorithms, which rely on associative operations, positioning DistPCA as a robust solution for the ever-growing demands of modern biobank-scale datasets.

The rest of this paper is organized as follows. In Section II, we present related efforts and discuss the motivation for this work. In Section III, we introduce our framework, providing a detailed analysis of the proposed parallelism scheme and the underlying numerical method. In Section IV, we present experimental results on the scalability of DistPCA on a multi-node HPC cluster and compare its performance with PCAone [36], the current state-of-the-art solution for out-of-core genomics PCA. Finally, in Section V, we summarize our key findings and discuss future directions.

## II. Background & Related Work

Given the individuals-by-SNPs (Single-Nucleotide Polymorphisms) normalized matrix, **A** ∈ ℝ^*m×n*^, genomic PCA aims to approximate the top-*k* PCs, that is, the *k* leading eigenvectors of the covariance matrix **AA**^*T*^ ∈ ℝ^*m×m*^. These eigenvectors are mathematically equivalent to the left singular vectors of **A** and are used interchangeably in this work.

Much of the prior work on genomic PCA has focused on optimizing the numerical method stage, a well-justified direction given that PCA boils down to computing the SVD, or equivalently the eigenvalue decomposition, of the covariance matrix associated with the normalized input data. For a covariance matrix of size *m × m*, extracting its eigenpairs requires 𝒪 (*m*^3^) operations, which becomes prohibitively expensive as the total number of individuals grows. Such limitations are addressed in Numerical Linear Algebra (NLA) using subspace projection schemes, which solve a much smaller eigenvalue problem by approximating the dominant eigenspace through a lower-dimensional subspace. While some early approaches adopted these schemes [27], [39], [40], they were predominantly in-core, assuming that the matrix could reside in main memory – an assumption that is untenable for modern largescale genomic datasets [35], [38].

To overcome this limitation, more recent methods adopt out-of-core designs that process data in blocks streamed from storage. One such method is FlashPCA2 [37], which relies on the Implicitly Restarted Arnoldi Method (IRAM) [41] and builds the approximation subspace through repeated passes over the data. Inspired by Randomized NLA (RandNLA), Bose *et al*. proposed TeraPCA [38], a framework that employs Randomized Subspace Iteration (RSI) and consistently outperforms FlashPCA2. Seeking to reduce the number of dataset passes required by prior approaches while maintaining comparable accuracy, Li *et al*. introduced PCAone [36], a framework built upon a window-based randomized SVD (RSVD) scheme, which was shown to outperform existing out-of-core genomic PCA methods.

Although these methods overcome the memory constraints of their in-core predecessors, they remain confined to single-node environments and rely primarily on optimizations provided by NLA multithreaded libraries (e.g., Intel MKL, Eigen, OpenBLAS). As a result, they do not fully address challenges related to large-scale data movement, I/O bottlenecks, and inter-node scalability, leading to degraded efficiency as genomic datasets continue to grow. To the best of our knowledge, DistPCA is the first out-of-core framework for genomic PCA that seeks to comprehensively accelerate all stages of the pipeline and bridges the gap between single-node and distributed execution, as shown in Table I. This gap is further underscored from the growing body of work adopting multi-node computing architectures for large-scale genomic pipelines [42], [43].

**TABLE I.** Parallelism in genomic PCA frameworks.

| Framework | I/O | Preprocessing | Numerical Method | Distributed |
| --- | --- | --- | --- | --- |
| SmartPCA [27] | ✗ | ✗ | ✓ | ✗ |
| FastPCA [39] | ✗ | ✗ | ✓ | ✗ |
| FlashPCA [40] | ✗ | ✗ | ✓ | ✗ |
| FlashPCA2 [37] | ✗ | ✗ | ✓ | ✗ |
| PCAone [36] | ✗ | ✓ | ✓ | ✗ |
| TeraPCA [38] | ✗ | ✗ | ✓ | ✗ |
| **DistPCA** | ✓ | ✓ | ✓ | ✓ |

## III. Methodology

**DistPCA** relies on a hybrid, multi-level, data parallelism scheme, inspired by the MapReduce paradigm [44], that combines multiprocessing, multithreading and SIMD vectorization to accelerate PCA in large-scale genomics. Built on top of MPI, the de facto message-passing standard for distributed memory programming, the framework supports out-of-core genomic PCA across compute nodes of a cluster, overcoming the scalability limitations of existing single-node approaches discussed earlier.

Parallelism is exploited through four distinct forms. First, at the process level, each MPI rank is assigned a disjoint subset of dataset blocks. Second, within each process, OpenMP threads are employed at block level to accelerate genotype normalization and Matrix-Multi-Vector (MMV) multiplications of RSI (thread-level parallelism). Third, genotype decoding is performed using SIMD vectorization, exploiting data-level parallelism within vector registers. Finally, a double buffering scheme is adopted within each process to overlap I/O operations with the computational pipeline, leveraging compute-transfer parallelism [45] to minimize resource idle time. Therefore, unlike the aforementioned frameworks (see Section II) that primarily focus on accelerating the numerical method stage of PCA, DistPCA adopts a more comprehensive approach by parallelizing not only the numerical method stage but also the two stages of the input data pipeline. Table I summarizes which stages of PCA are parallelized by each framework, and whether they support multi-node (distributed) execution.

Below, a brief overview of the DistPCA framework is presented, describing the numerical backbone along with the multi-level hybrid parallelism scheme for out-of-core computation of the sought PCs.

### A. Randomized Subspace Iteration

At its core, DistPCA employs the RSI method, as described in the supplementary material of [38], to compute the leading PCs. Briefly, RSI is a multivector generalization of the power method to extract multiple eigenpairs. It approximates the invariant subspace spanned by the *k* leading eigenvectors (with *k* ≪*m*) of the *m* × *m* covariance matrix **AA**^*T*^ of the normalized genotype matrix. In practice, to obtain accurate approximations of the leading *k* eigenvectors, RSI targets a slightly larger subspace of dimension *s* ≥ *k*, typically chosen as a modest oversampling of *k* (often on the order of 2*k*).

The method, outlined in Algorithm 1, initializes the target subspace with a random Gaussian matrix **X**_0_ ∈ ℝ^*m×s*^ (Step 1) and iteratively refines it to approximate the dominant eigenspace of **AA**^*T*^ via *p* power iterations (Steps 3–9). To ensure numerical stability, periodic orthogonalization is incorporated (Steps 5 and 8), followed by subspace projection to form a smaller size matrix **M** (Step 10). Solving the smaller eigenvalue problem, defined by **M**, provides an approximation to the top-*k* eigenpairs (Steps 11 and 12). This process is repeated until convergence is achieved, eventually yielding the *k* leading left singular vectors of **A**. Notably, each iteration requires *p* + 1 passes over the dataset. However, empirical evaluation in [38] demonstrated that a single application of the covariance matrix is often sufficient to recover the top 10–20 PCs with satisfactory accuracy, thereby reducing the number of passes over the dataset to two per iteration.

#### Algorithm 1

Randomized Subspace Iteration

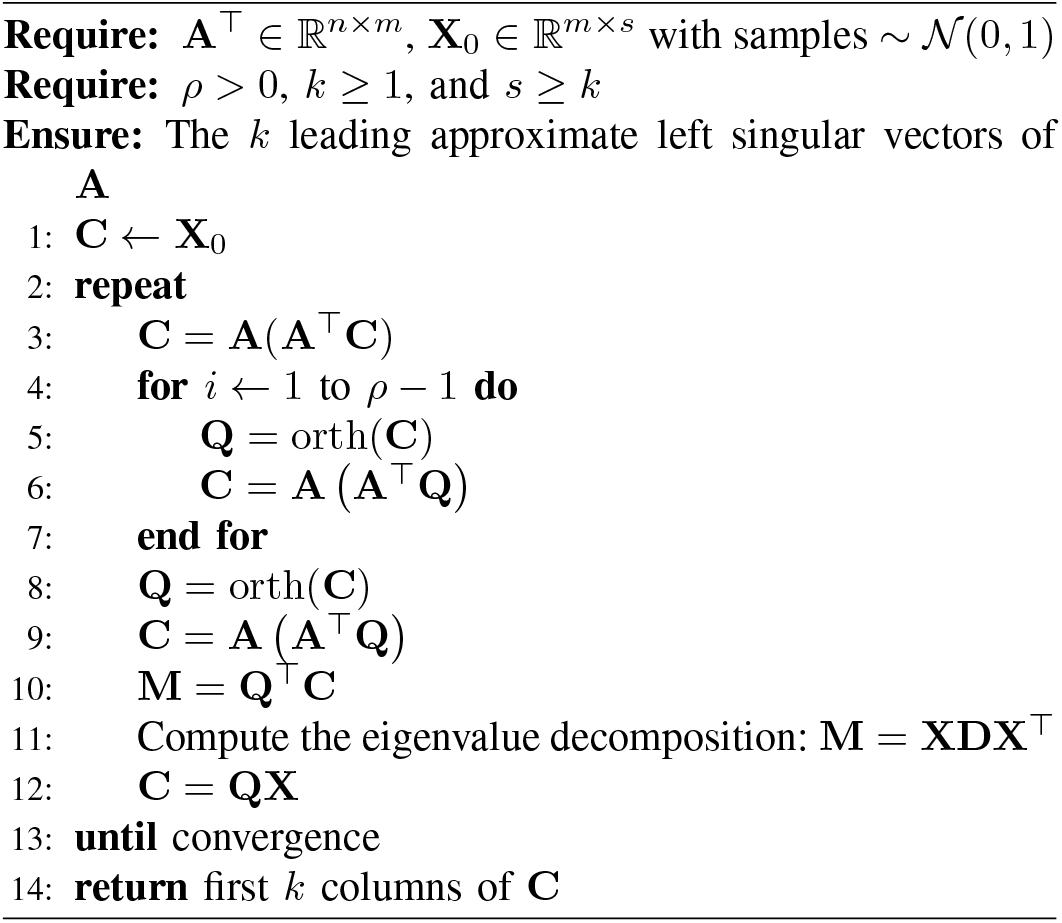

#### Algorithm 2

Out-of-Core MMV: **C** = **A**(**A**^*T*^**X**)

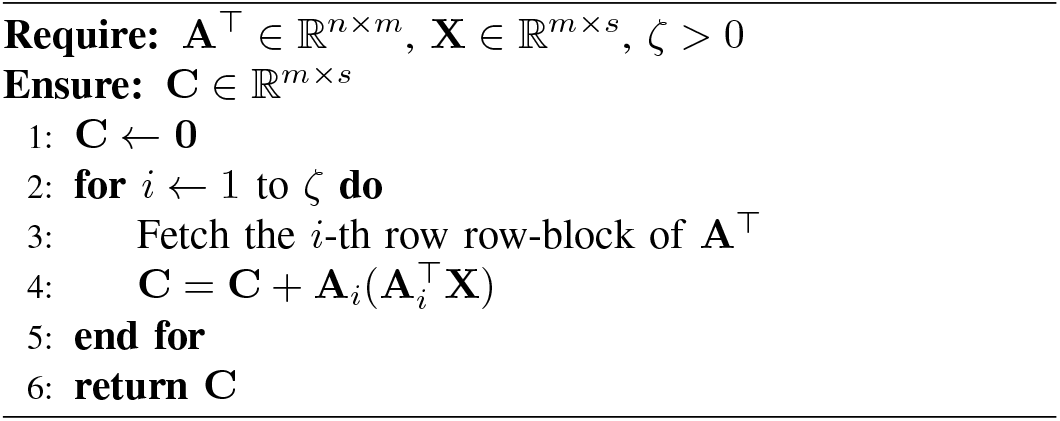

From an implementation perspective, the RSI algorithm is well suited for out-of-core PCA, as it relies on repeated MMV multiplications with the covariance matrix. As listed in Algorithm 2, these operations can be decomposed into independent contributions from subsets of rows of **A**, enabling incremental processing of the dataset in blocks loaded from storage. In addition, when built on top of Intel MKL, GNU Scientific Library (GSL) or other optimized NLA libraries, the efficiency of RSI can be further improved utilizing multithreaded BLAS3 operations. For a more detailed description of the RSI algorithm, we refer the reader to [38].

### B. Hybrid Multi-Level Parallelism Scheme

Figure 2 provides an overview of the proposed hybrid multilevel parallelism scheme. Initially, the raw data file, stored as a PLINK .*bed* file in SNP-major format, is logically divided into *P* chunks, where *P* corresponds to the number of MPI processes. Each chunk consists of a sequence of consecutive, equally-sized, non-overlapping blocks and is assigned exclusively to a single MPI rank. For each chunk, the final block may contain fewer samples if the total number of genotypes is not evenly divisible by the selected block size.

**Fig. 1.**
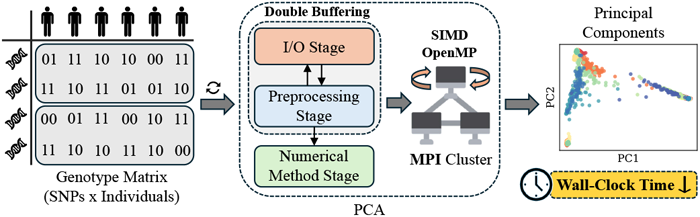
Overview of the DistPCA framework. The input genotype matrix (left) is stored in SNP-major format. PCA is performed through a block-aware three-stage pipeline (center) accelerated via a multi-level parallelism scheme that leverages MPI, OpenMP, and SIMD vectorization, with double-buffered I/O, collectively reducing wall-clock time of PCs computation (right).

**Fig. 2.**
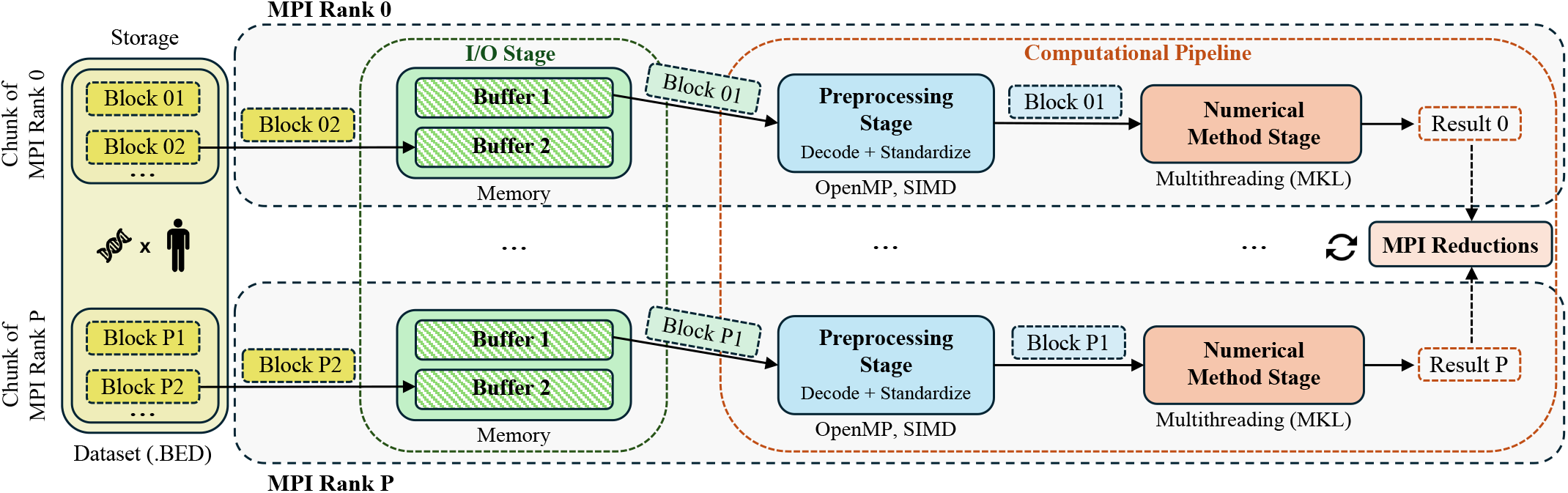
Overview of DistPCA multi-level hybrid parallelism scheme. The input dataset is partitioned across MPI ranks, which fetch blocks in parallel using double-buffered I/O, preprocess them via OpenMP/SIMD, and perform multithreaded MMV computations, followed by MPI reduction operations to update the global eigenspace.

At each iteration of RSI, all MPI processes concurrently fetch and process their assigned data blocks. Therefore, at any given time *t*, at most *P* blocks are simultaneously loaded into memory for processing. Within each process, for each block, once the genotype samples are loaded and decoded via SIMD vectorization using SSE4.2 intrinsics, standardization is applied in parallel using OpenMP threads. The block’s contribution to the current subspace approximation is then computed through multithreaded BLAS-3 MMV multiplications, according to Algorithm 2. After completing the local computations for all assigned blocks, MPI processes aggregate their partial results through MPI (blocking) collective communication, yielding the updated eigenspace approximation for the entire dataset, which is subsequently broadcast to all ranks.

As presented in Algorithm 1, in each iteration, MPI processes fetch their assigned blocks from storage twice, once for each MMV multiplication with the covariance matrix, and perform two collective communications to aggregate their partial results into matrix **C**. When power iterations are applied (i.e., *ρ >* 1), each additional iteration entails an extra pass over the data blocks and an additional collective communication for the update of matrix **C**. For data normalization, the mean and standard deviation of each sample must be computed beforehand. Since these statistics have a small memory footprint, their computation is performed during the initial block fetch for each sample and cached for reuse in subsequent iterations, thereby avoiding redundant computations.

Parallel block fetching is enabled by using binary offsets that indicate the starting position of each block within the input file. This design allows each MPI process to move (seek) directly to its assigned blocks and read them independently, without relying on incremental file parsing. Although this parallel MPI I/O approach significantly improves throughput, data transfers from secondary memory may still incur significant latency, leading to CPU idle periods while waiting for I/O completion. To mitigate this effect and hide the latency of data transfers between storage and main memory, each MPI process employs a double-buffering scheme in which I/O and computation proceed concurrently [46]. Specifically, two data buffers are used per process (see Figure 2): while data preprocessing and numerical computations of RSI are performed on a block stored in the first buffer, the next block is simultaneously read from storage into the second buffer. By overlapping computation with I/O, this scheme minimizes idle time and ensures a steady data flow to the available CPU cores.

A key parameter of the proposed scheme is the block size used to partition the dataset, as this determines the number of SNPs processed by each MPI rank at a time and, thus, the parallel I/O performance. The optimal size depends on several factors, including dataset size, RAM, I/O bandwidth, Last-Level Cache (LLC), and the number of compute nodes. In practice, the block size should be selected such that the I/O workload generated by the workers matches the storage system’s capabilities (e.g., bandwidth and latency) and maximizes the LLC hit ratio. Such an approach prevents cache contention and memory hierarchy pressure, resulting in high throughput with low latency. Consequently, the processing rate (samples per second) increases, improving the runtime performance.

## IV. Evaluation

The performance of DistPCA was evaluated on three synthetic datasets and three publicly available real-world datasets of varying sizes, as summarized in Table II. Throughout all experiments, RSI targets the leading *k* := 20 PCs, starting from an initial approximation subspace of dimension 2*k*, with a fixed block size of 100 SNPs. Consistent with prior studies [36], [39], convergence is determined via the mean explained variance (MEV) of eigenvectors, a metric for evaluating the quality of estimated PCs, and iterations are terminated once the relative change in MEV falls below a threshold of 10^*−*3^. The implementation of DistPCA is publicly available at Open Science Framework (OSF) ^1^.

**TABLE II.** Evaluation Datasets.

| Name | Individuals | SNPs | Type | Size (.BED file) |
| --- | --- | --- | --- | --- |
| SGDP | 345 | 694,659 | Real | 58 MB |
| HGDP | 942 | 133,594 | Real | 31 MB |
| 1000 Genomes | 2,490 | 1,664,505 | Real | 989 MB |
| 50K Genomes | 50,000 | 6,000,000 | Synthetic | 75 GB |
| 500K Genomes | 500,000 | 3,000,000 | Synthetic | 350 GB |
| 1M Genomes | 1,000,000 | 1,000,000 | Synthetic | 233 GB |

To assess the scalability of the proposed hybrid parallelism scheme, we performed multiple runs with different worker configurations (i.e., number of MPI ranks), while keeping the number of OpenMP threads per rank fixed at 8, unless otherwise stated. Following prior work on performance evaluation [47], [48], the reported execution times correspond to the median of 5 benchmark runs for each configuration. Beyond runtime evaluation, we also conducted an accuracy study on the *1000 Genomes* dataset, which is small enough to perform full-rank SVD, by computing the relative error between the 10 leading eigenvectors estimated by our framework and those from full-rank SVD. Since the reliability of the RSI method has been thoroughly validated in [38], this brief analysis is primarily included to confirm that our parallelism scheme adequately preserves the data dependencies inherent to the numerical method, and, thus, does not affect the quality of the *k*-leading computed PCs.

### A. Experimental Setup

#### Datasets

As mentioned earlier, experiments were conducted on six distinct datasets (Table II). Across these datasets, the number of individuals ranges from 345 to 1M, while the number of SNPs varies from 130K to 6M. Although stored compactly in PLINK binary format, several of these datasets expand to multiple terabytes in uncompressed double-precision format (reaching up to 11 TB), enabling the evaluation of the proposed scheme in out-of-core scenarios. For the real datasets, a standard quality control (QC) step was applied using the PLINK toolset [34]. Variants with minor allele frequency (MAF) below 0.01 were removed, followed by a linkage disequilibrium (LD) pruning with a 1,000 KB window size, a variance inflation factor of 50 SNPs, and an *r*^2^ threshold of 0.2. The synthetic datasets were generated using the Pritchard-Stephens-Donnelly (PSD) model for genotype simulation [49], [50]. Individual ancestry proportions were drawn from the PSD model fitted to the *1000 Genomes* dataset, while per-population allele frequencies were simulated using Wright’s *F*_*ST*_ and the Weir and Cockerham estimator [51].

#### Hardware Configuration

All experiments were conducted on the ARIS supercomputer, a national HPC cluster facility, using four thin compute nodes. Each node is equipped with a dual-socket AMD EPYC 7742 CPU (128 cores, 2.25 GHz), partitioned into eight Non-Uniform Memory Access (NUMA) domains (four per socket), with 512 GB of RAM. Within each socket, each NUMA node comprises two Core Complex Dies (CCDs), each featuring eight CPU cores that share a 32 MB LLC (L3). On top of that, ARIS provides high-performance storage through IBM’s GPFS. For each worker configuration, MPI ranks were distributed across NUMA domains, while OpenMP threads were pinned to the cores within each domain to improve memory locality and prevent thread migration. For all experiments, hyperthreading was disabled, while the available physical memory per node was restricted to 64 GB, regardless of the total system memory. MKL routines were accessed via Intel oneAPI (v2025.0.1).

#### Baseline

As discussed in Section II, prior research had primarily focused on accelerating the numerical stage of PCA using single-level shared-memory parallelism (i.e., multithreading), without support for distributed execution. Hence, a direct one-to-one performance comparison with existing frameworks is not straightforward, as our approach is designed for multi-node cluster environments and employs a hybrid multi-level parallelism scheme that targets not only the numerical stage but also the two stages of the input data pipeline. Nevertheless, to provide a comprehensive evaluation of DistPCA, we conducted a comparative study of its runtime performance against PCAone, which has been reported in [36] to consistently outperform all other frameworks discussed in Section II. Our objectives in the present study are twofold: (i) to confirm, as reported in prior studies [36], [38], that in tera-scale genomics PCA, repeated I/O operations for fetching blocks of data from storage create a significant performance bottleneck, and (ii) to demonstrate that accelerating the input data pipeline via the proposed scheme results in clear performance improvements over existing frameworks.

### B. Results

We first evaluate the scalability of DistPCA focusing on the synthetic datasets and the *1000 Genomes* dataset. As shown in Figure 3, the proposed MPI-based parallelism scheme demonstrates strong scalability, achieving near-linear speedup and maintaining parallel efficiency above 82% across all evaluated scenarios.

**Fig. 3.**
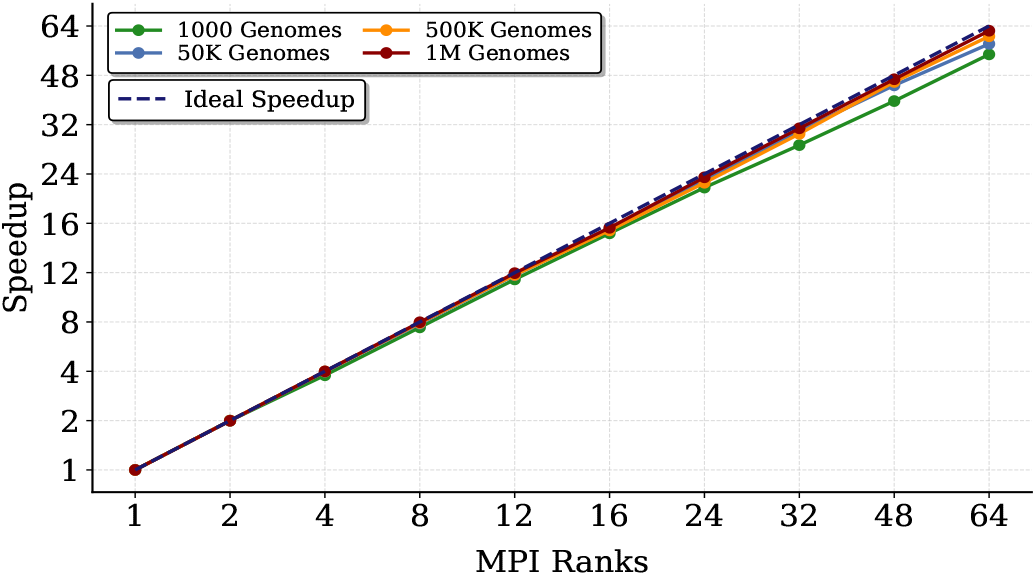
Strong scaling speedup of DistPCA. Across all datasets and worker configurations, parallel efficiency remains above 82%.

Regarding the *50K Genomes* dataset, increasing the number of MPI ranks from 1 to 64 results in a wall-clock time reduction from 61 hours to just over one hour, which corresponds to 58.2× speedup and more than 98% time improvement, as depicted in Figure 4. Similarly, for the two other synthetic datasets, whose uncompressed size exceeds 7 TB, using 32 MPI ranks reduces the computation time of the first 20 PCs to under 40 minutes, yielding a time reduction exceeding 96%. The wall-clock time gap of these two datasets compared to the *50K Genomes* dataset can be primarily attributed to: (i) the lower number of RSI iterations required for convergence (20 vs. 89), and (ii) their smaller number of SNPs, resulting in fewer data block fetches and, thus, reduced I/O overhead. As for the *1000 Genomes* dataset, configurations beyond 24 MPI ranks exhibit lower parallel efficiency compared to the synthetic datasets. This is due to its relatively smaller size (in uncompressed format *≈* 33 GB), which, combined with Amdahl’s law and the imposed communication overhead, limits the attainable speedup. Despite this, parallel efficiency remains above 82% across all configurations, allowing for a maximum achieved speedup of 54.8× (64 MPI processes). Results from the *SGDP* and *HGDP* datasets are omitted from the performance graphs, since their small size leads to the computation of the 20 leading PCs in only a few seconds, even when using 8 MPI processes. Nevertheless, DistPCA maintains parallel efficiency above 94% on both datasets. Overall, the proposed framework demonstrates strong scalability with negligible communication overhead, achieving near-linear speedup in a multi-node cluster and time improvements of up to 98.4%.

**Fig. 4.**
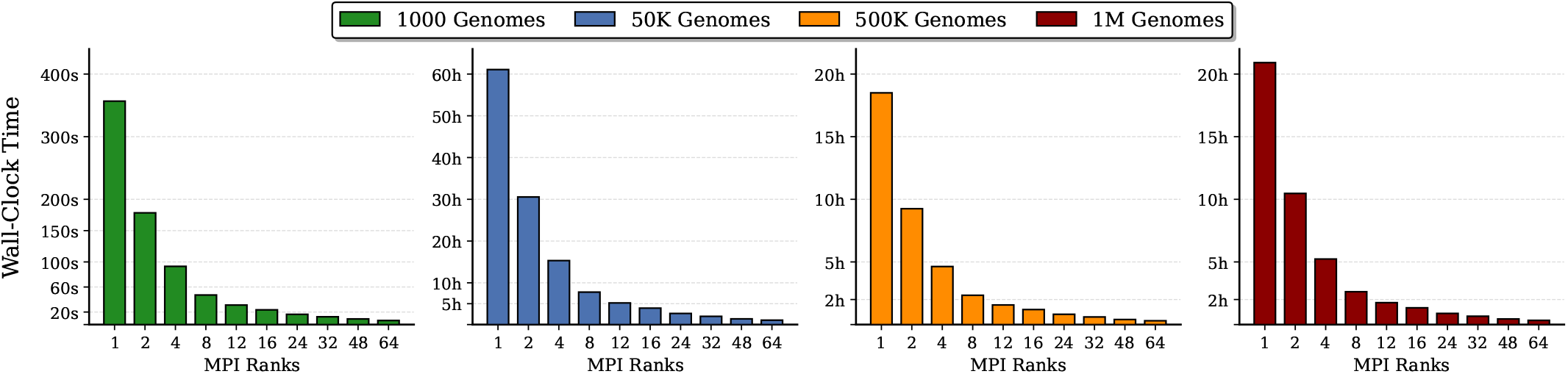
Runtime performance of DistPCA on the *1000, 50K, 500K*, and *1M Genomes* datasets. As observed, for datasets exceeding 70 GB in .*bed* format, computing PCs with a single MPI rank requires at least 17 hours. Across all datasets, increasing the number of MPI ranks from 1 to 64 results in a wall-clock time reduction of over 98%. Results from the *SGDP* and *HGDP* datasets are omitted, as their runtimes remain below 5 seconds even with 8 MPI ranks.

Table III presents the wall-clock times achieved by DistPCA and PCAone [36] when applied to the synthetic datasets and the *1000 Genomes* dataset. To ensure the comparison was as fair as possible, the total number of workers was fixed to 64 across all experiments, regardless of whether parallelism was applied at the thread level or through a hybrid scheme (i.e., MPI processes × OpenMP threads). Accordingly, PCAone was employed using 64 OpenMP threads, while DistPCA was configured with 8 MPI processes and 8 OpenMP threads per process.

**TABLE III.** Performance Comparison with PCAone.

| Dataset | PCAone | DistPCA | Speedup | Reduction % |
| --- | --- | --- | --- | --- |
| 1000 Genomes | 173s | **47s** | 3.68x | 72.8% |
| 50K Genomes | 9.1h | **7.8h** | 1.17x | 14.1% |
| 500K Genomes | 12.1h | **2.3h** | 5.26x | 78.5% |
| 1M Genomes | 7.9h | **2.6h** | 3.04x | 67.9% |

As observed from Table III, DistPCA consistently outperforms PCAone across all datasets. In particular, for the *1000, 500K*, and *1M Genomes* datasets, DistPCA achieves execution times that are at least 3× faster than PCAone. For the *50K Genomes* dataset, the performance gap narrows, as PCAone converges in 7 iterations, whereas the RSI method requires 89 iterations. Nevertheless, DistPCA still provides a time reduction of 14.1%. It is important to note that PCAone is confined to single-node execution and thus scales only with the resources of a single machine, whereas DistPCA is designed for distributed environments and can continue to scale with additional compute nodes.

Our results are consistent with prior studies indicating that PCA runtime is dominated by I/O overhead [36], [38], which limits the attainable speedup when parallelism is applied only to preprocessing and/or algebraic operations, as in PCAone. By contrast, the proposed framework employs a multi-level hybrid parallelism scheme that targets all stages of PCA, resulting in improved scalability and time reductions of up to 78.5%.

Regarding the accuracy of estimated PCs, Figure 5 reports the entry-wise relative error of the 10 leading eigenvectors computed by DistPCA for the *1000 Genomes* dataset, compared to those obtained via full-rank SVD. We observe that the relative error is approximately uniform across the entries of each eigenvector. As expected, eigenvectors associated with the largest eigenvalues are captured more accurately due to faster convergence. Overall, the observed error is consistent with that reported in TeraPCA [38], which also employs the RSI method for PCs estimation. Therefore, the parallelism scheme of our framework adequately preserves RSI data dependencies and does not affect the quality of the computed PCs. Beyond entry-wise accuracy, it is important to verify that the estimated PCs retain their ability to capture meaningful population structure. To this end, Figure 6 shows the projection of individuals of the *1000 Genomes* dataset on the first two estimated PCs, colored by population. The resulting structure demonstrates clear population clustering consistent with known stratification patterns [38], [52].

**Fig. 5.**
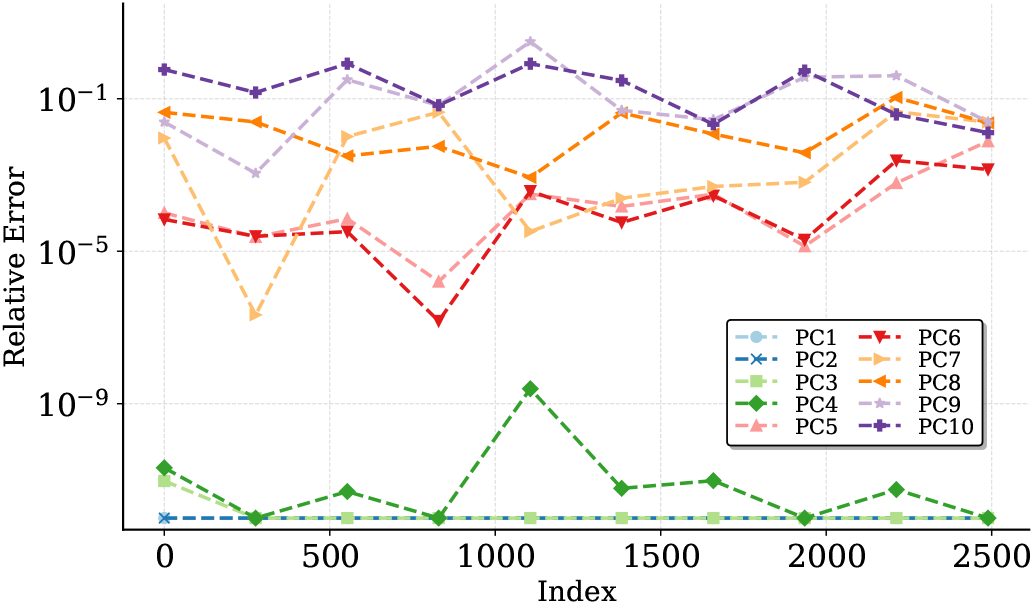
Entry-wise relative error of the 10 leading eigenvectors computed by DistPCA for the *1000 Genomes* dataset, compared to the eigenvectors returned by the full-rank SVD. Each eigenvector has 2,491 entries.

**Fig. 6.**
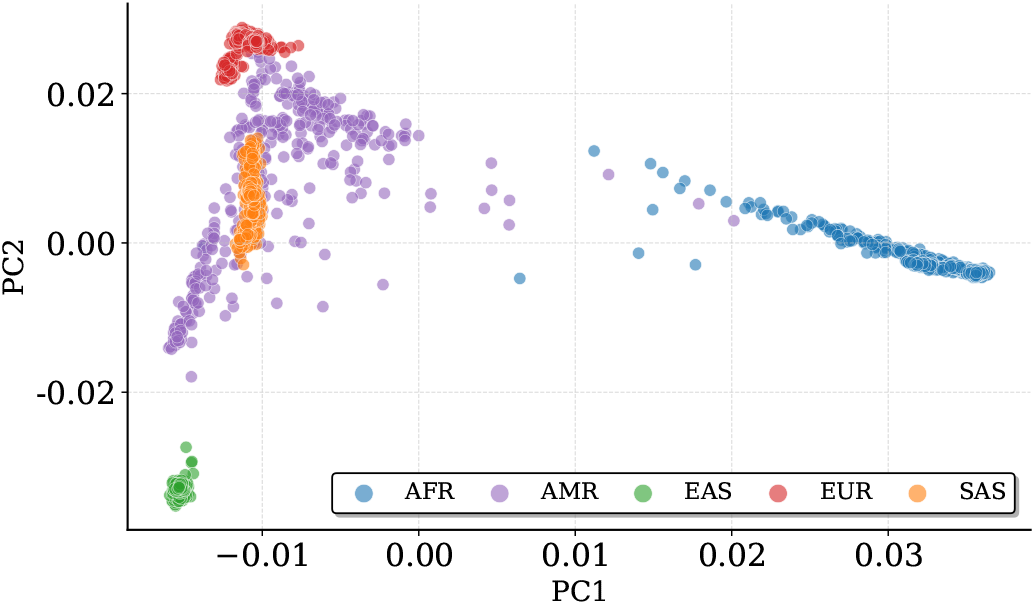
Projection of the samples of the *1000 Genomes* dataset on the top two left singular vectors (PC1 and PC2), as computed by DistPCA. Samples are grouped into five populations: AFR (African), AMR (Ad Mixed American), EAS (East Asian), EUR (European), and SAS (South Asian).

## V. Conclusion

We introduced DistPCA, a distributed C++ framework for large-scale genomic PCA that efficiently combines MPI-based distributed execution with OpenMP multithreading, SIMD vectorization, and compute-transfer overlap via double buffering to comprehensively accelerate the three key stages of PCA. Experimental results on real and synthetic datasets demonstrate strong scalability, near-linear speedups and reduction in wall-clock time by up to 98%, while maintaining the accuracy of the recovered PCs and outperforming existing frameworks. These results highlight the importance of codesigning data movement (I/O) and computation in genomic PCA, establishing DistPCA as a robust and scalable solution for the growing demands of modern biobank-scale datasets.

In future work, we aim to extend DistPCA to support alternative block-based numerical methods, broadening its applicability across different algorithmic schemes. Moreover, we plan to integrate heterogeneous hardware architectures, particularly GPUs, to further accelerate the computational parts of PCA and achieve additional performance gains on modern HPC infrastructures.

## VI. Acknowledgments

The authors would like to thank Professor Petros Drineas for his valuable advice and insightful feedback. They also gratefully acknowledge the Computer Center of the Department of Computer Engineering and Informatics at the University of Patras, Greece, for providing the computational resources used during the implementation and early stages of DistPCA. This work was supported by computational time granted by the National Infrastructures for Research and Technology S.A. (GRNET S.A.) at the National HPC facility ARIS under project ID pa260203distpca.

## Footnotes

1 https://osf.io/63sqh/overview?view_only=2d26f7cfb3bc4d36954775f6219992a2

